# Dual-function logic gates based on CRISPRi

**DOI:** 10.1101/2024.10.23.619786

**Authors:** Zhiwu Yao, Shaobin Guo

## Abstract

Developing logic gate circuits that are programmable and reliable for specific functions is a central goal in synthetic biology. Traditional synthetic circuits often rely on protein regulators, which face scalability and resource burden limitations. To overcome these challenges, we introduce a novel approach utilizing dual-function and multi-level logic gates based on the CRISPRi system with dCas9 and sgRNA. This method, implemented in an in vitro transcription system, enables the rapid design and validation of complex logic gates. Our dual-function and multi-level design strategies achieve robust functionality in NOR, NAND, AND, and OR gates, allowing different logic outputs by altering only the inputs while reusing all other modules. This showcases significant advancements in efficiency and scalability for synthetic circuit construction. This work reduces development time and simplifies circuit design, paving the way for more efficient synthetic biology applications.

## Introduction

Constructing programmable, predictable artificial decision circuits with specific functions is one of the main focuses of synthetic biology^1–3^. Since the emergence of synthetic biology in 2000, most synthetic circuits have relied on protein regulators (mostly transcription factors) for control^4–6^. As researchers build more complex synthetic circuits, it has become apparent that transcription factors have low scalability; each new protein must be retested for orthogonality with the remaining proteins in the circuit, making it difficult to construct more complex gene circuits using transcription factors^7^. Additionally, the high expression levels of various proteins consume significant cellular resources^8^.This increased burden on chassis cells can lead to slower growth, reduced metabolic performance, and evolutionary instability^8^.

Recently, there has been growing interest in RNA regulator-based circuits to overcome the limitations of protein regulators^9^. Among them, the prokaryotic adaptive immune system CRISPR provides a powerful platform for constructing RNA-driven synthetic circuits^10–12^. Catalytically inactive mutants like dCas9 can be easily directed to nearly any sequence via single guide RNA molecules (sgRNA)^13^. When targeted to the prokaryotic promoter (or downstream) region, the spatial hindrance of the dCas9-sgRNA complex results in transcriptional repression, a method known as CRISPR interference (CRISPRi)^14^. CRISPRi presents several advantages in synthetic circuit design compared to protein regulators. Due to its RNA-guided nature, CRISPRi is highly programmable^15,16^, allowing for the easy design of highly orthogonal sgRNAs^17,18^ and straightforward prediction of behavior in different environments through computer simulations^19,20^.Integrating proteins of different functions to the dCas9 protein further broadens the dCas9 protein lineage, providing a promising toolkit for constructing functional and efficient gene circuits^21–26^. Moreover, CRISPRi imposes a lower burden on host cells, as its coding sequence is shorter than that of protein-based repressors, simplifying circuit manipulation and delivery^27–29^.

However, the overexpression of dCas9 protein within cells can still inhibit normal cell growth and cause cell abnormalities^13,30,31^. Therefore, our work focuses on prototyping and developing the CRISPRi system in a cell-free system. Cell-free systems (CFS) have become an attractive platform for gene circuit prototyping, synthetic cell construction, and engineering biosynthetic pathways^32– 36^. However, there are few examples of gene circuits capable of dynamic, multi-gene regulation in CFS^17,37^.After thoroughly studying the principles of CRISPRi repression, we chose the T7 RNAP system as a foundation for further developing a CRISPRi in vitro system. Unlike CFS, the T7 RNAP system involves only transcription and its associated metabolic components, characterized by typically low protein and RNA turnover rates. Researchers have already constructed and implemented multi-level coupled riboswitches in the T7 RNAP system^38,39^. Since CRISPRi inhibits gene expression at the transcriptional level, we can develop a CRISPRi system with the T7 RNAP system.

A significant bottleneck in the rapid prototyping of biological circuits is the need to independently design new input, output, and information processing modules for each logic gate, drastically increasing the workload. Inspired by design concepts from electronic circuits, we propose a new design approach based on CRISPRi. In this approach, two or more logic gates differentiate only at the input module, while their output and information processing modules are identical. By merely altering the input, the output can be correspondingly changed. Based on this concept, we designed NOR and NAND gates with identical information processing and output modules, requiring only different sgRNAs to achieve distinct logical outputs. After further validating that sgRNAs can serve as multi-level cascading signals, we designed and validated multi-level AND and OR gates with the same information processing and output modules. We believe our streamlined approach provides a valuable design principle which not only simplifies the design process but also significantly reduces the time and resources required for the development of complex biological circuits.

## Materials and methods

### Plasmid Construction

All sequences and plasmids used in this study are listed in Supplementary Tables S1. All DNA oligonucleotides were purchased from Fuzhou Sunya Biotechnology. The plasmid pBR322-backbone contains mainly functional fragments of the 3WJdB fluorescent aptamer sequence driven by the T7 promoter based on the design of the pBR322. pBR322 contains a carbenicillin resistance gene and an origin of replication ColE1 to allow its passage through *E. coli*. All the gene fragments (Fuzhou Sunya Biotechnology) were ligated using Golden Gate assembly (#M0551S, New England Biolabs) and the ligated product was transformed into *E. coli* DH5α strain and cultured on LB agar plate with appropriate antibiotics (100 μg/mL carbenicillin, Shanghai yuanye Bio-Technology). Single colonies were inoculated in LB liquid medium with appropriate antibiotics, and cultured for 14 h at 37°C, 220 rpm. A volume of 1 mL of cultured bacteria was taken for sequencing (Fuzhou Sunya Biotechnology). The bacterial strains with correct sequencing results were preserved, and the plasmids were extracted

### In Vitro Transcription Reaction and Fluorescence Assays

To prepare linearized templates for in vitro transcription, the plasmids were used as templates for polymerase chain reaction (PCR) with primers to amplify regions including T7 promoter and desired transcript (#DC301-01, Vazyme Biotech). In vitro transcription reactions were prepared on ice with 2-50 nM of linearized DNA template, 40 μM of DFHBI-1T (#SML2697, Sigma-Aldrich), 0.5 mM of NTPs, 1.5 μL of T7 RNAP (50 U/μL), and 0.75 μL of ribonuclease inhibitor (#R301-03, Vazyme Biotech) in the reaction buffer (50 mM Tris-HCl pH 7.9, 10 mM MgCl_2_, 100 mM NaCl, 100 μg/ml Recombinant Albumin) (#B6003V, New England Biolabs) at a final volume of 30 μL. The in vitro transcription reaction was conducted in a 384-well plate (Agilent) with three replicates per experimental condition, including a control reaction where the DNA template was excluded. The reaction temperature was controlled at 37°C and the fluorescence signal to determine the transcription level of 3WJdB was measured using a plate reader (Molecular Devices). The 3WJdB fluorescence (excitation/emission 472/507 nm) was taken 2 hours after the start of the in vitro transcription reaction.

### Computational Design of Orthogonal Repressors

Randomly generated 23bp DNA fragments were generated using a program of our own design, see Appendix for specific program files. The randomly generated fragments were submitted to the website http://crispor.tefor.net/, the target genome was selected as Escherichia coli BL21(DE3)-NCBI GCA_000022665.2(ASM2266v1), and the PAM site was selected as 20 bp NGG-Sp Cas9. According to the final returned results, the high scoring, multi-APM site sequences were selected, and these sequences were excerpted in a table, and subsequently a program was designed to calculate the Hamming distance between each sequence, and the two with the furthest Hamming distances were selected as a set of random orthogonal fragments. The specific program is shown in the Appendix.

### sgRNA Design and Cloning

All the sgRNAs used in this work were designed via the crispor (at http://crispor.tefor.net/), setting a guide length of 20 nucleotides, GCA_000022665.2 as reference genome, and NGG was used as the PAM site for the target sequence.

## Results and discussion

### (1) T7 RNAP system mimics cellular transcription

T7 RNAP exhibits high specificity for the T7 promoter in vitro, effectively simulating the transcription process within cells. The buffering environment provided by the T7 RNAP system is well-suited for the optimal functioning of T7 RNAP; however, it may not necessarily be conducive to the functionality of the dCas9 protein. To develop a CRISPRi system with the T7 RNAP framework, we first combined and integrated the T7 RNAP buffer with the dCas9 protein buffer, ensuring that both proteins can perform their functions effectively (data not shown).

Next, we selected fluorescent RNA aptamers (FLAPs) as reporter genes^40,41^. When bound to fluorescent dyes, these aptamers can produce fluorescence intensity comparable to that of fluorescent proteins, making them an excellent substitute for fluorescent proteins in cell-free environments. Besides, with FLAPs, we can investigate only the transcription process without the complication from translation as CRISPRi mainly affects transcription.

Subsequently, we selected two different fluorescent aptamers as output signals for further validation. First, we verified the effect of varying concentrations of linear DNAs containing these aptamers during transcription on the resulting fluorescence intensity, as depicted in Fig. 1a-b. With an increase in the linear template DNA strand concentration during transcription, the fluorescence intensity was correspondingly enhanced.

**Figure 1:**
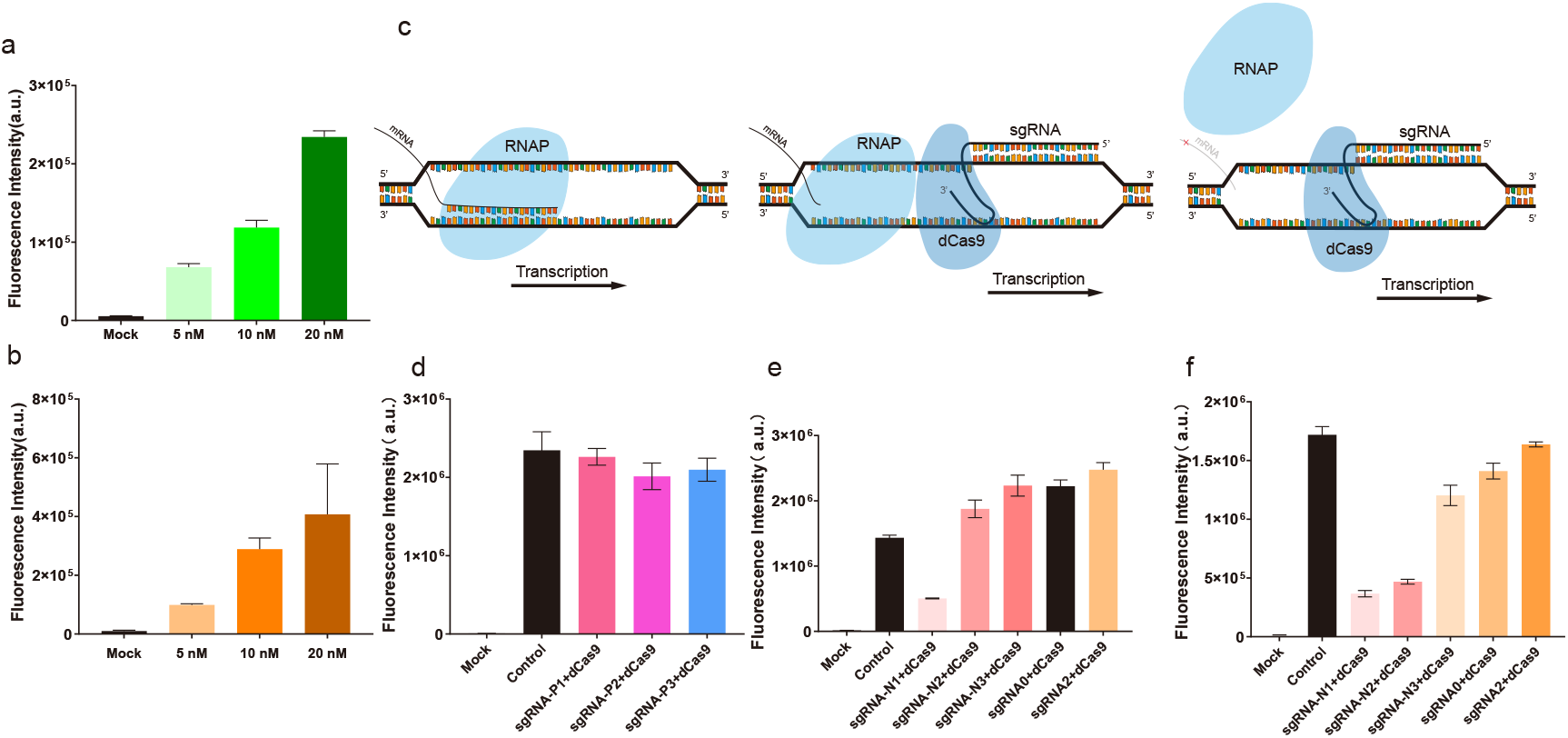
Fluorescence values of 3WJdB aptamer transcription at different concentrations. b: Fluorescence values of 4Pepper aptamer transcription at different concentrations. c: Schematic representation of CRISPRi-mediated transcriptional repression, where the sgRNA-dCas9 complex exerts steric hindrance to block RNA polymerase (RNAP) elongation, thereby inhibiting transcription. d: Inhibitory effects of various sgRNAs designed on the 4Pepper fluorescent aptamer. 1e: Inhibitory effects of various sgRNAs designed on the 3WJdB fluorescent aptamer, differing from Figure 1f in that the sgRNAs were pre-transcribed and purified in advance. f: Inhibitory effects of co-transcription of sgRNAs and the 3WJdB fluorescent aptamer. Error bars represent the standard deviation (s.d.) of three biological replicates.

### (2) CRISPRi in vitro validation

CRISPRi mainly consists of dCas9 protein and sgRNA. The repression of transcription is caused by sgRNA-dCas9 complex exerting steric hindrance to block RNA polymerase (RNAP) elongation (Fig. 1c). We employed sgRNA as a delivery signal to further construct a multi-level cascade of higher-order gene circuits, conceptualizing each level of CRISPRi as a node representing a discrete transcription unit. To characterize individual CRISPRi nodes, we constructed separate plasmids or linear DNA fragments for each of the dCas9 proteins, sgRNAs, and target sequences, thereby enabling expression level-independent titration of all CRISPRi components.

During the individual node CRISPRi characterization, we designed sgRNA targeting sites for the two aforementioned fluorescent aptamers, with three sites designated for each aptamer. Subsequently, the dCas9 protein was co-incubated with the sgRNA at 37°C for 20 minutes before being mixed with the T7 RNAP system containing the fluorescent aptamer. The strength of the CRISPRi inhibition effect was then determined by measuring the fluorescence values (Fig. 1d-e). Our observations revealed that increasing the linear expression concentration of sgRNA resulted in higher levels of repression (data not shown).

As shown in Fig. 1e, we successfully verified the feasibility of CRISPRi in the T7 RNAP in vitro transcription system. Among the three sgRNA sites designed for 3WJdB, only sgRNA-N1 demonstrated a significant inhibitory effect. Fig. 1d illustrates the inhibitory effect of the three sgRNA sites of 4Pepper, with only sgRNA-P3 exhibiting little inhibitory effect. This can be explained by the strong secondary structure formed by 4Pepper, which hindered the targeting of specific sequences by the dCas9 protein-sgRNA complex. As a result, 3WJdB was chosen as the output gene for the following experiments.

Noted that in Fig. 1e, the fluorescence intensity of the experimental group is greater than that of the positive control group. This can be attributed to the fact that sgRNA is transcribed and purified using the T7 RNAP system. During the purification process, the majority of enzymes and DNA strands are removed; however, a fraction of small molecules, such as NTPs, remain. As a result, the NTPs are introduced alongside the sgRNA, leading to a higher resource concentration in the T7 RNAP system of the experimental group compared to the positive control group.

To address the issue of the fluorescence signal in the experimental group being higher than that of the positive control group in subsequent studies, we considered that both sgRNA and 3WJdB utilize the T7 promoter and the same system. Therefore, it is theoretically feasible to transcribe them simultaneously within the same system. The reason we initially avoided this approach was that if sgRNA and 3WJdB are transcribed simultaneously, 3WJdB will begin transcription and generate fluorescent signals while sgRNA has just begun to be transcribed, resulting in insufficient sgRNA to bind to dCas9 and inhibit 3WJdB. This would lead to a decrease in the inhibitory efficiency of CRISPRi. However, the results of co-transcription showed that, similar to the transcription of sgRNA alone, CRISPRi exhibited inhibition after 10 minutes, without significant delay in the inhibitory effect (Fig. 2a). Analyzing the inhibitory effect of co-transcription, we found that the inhibitory effect was enhanced, with inhibition being effectively doubled with sgRNA-N1 and sgRNA-N2 (Fig. 1f).

**Figure 2:**
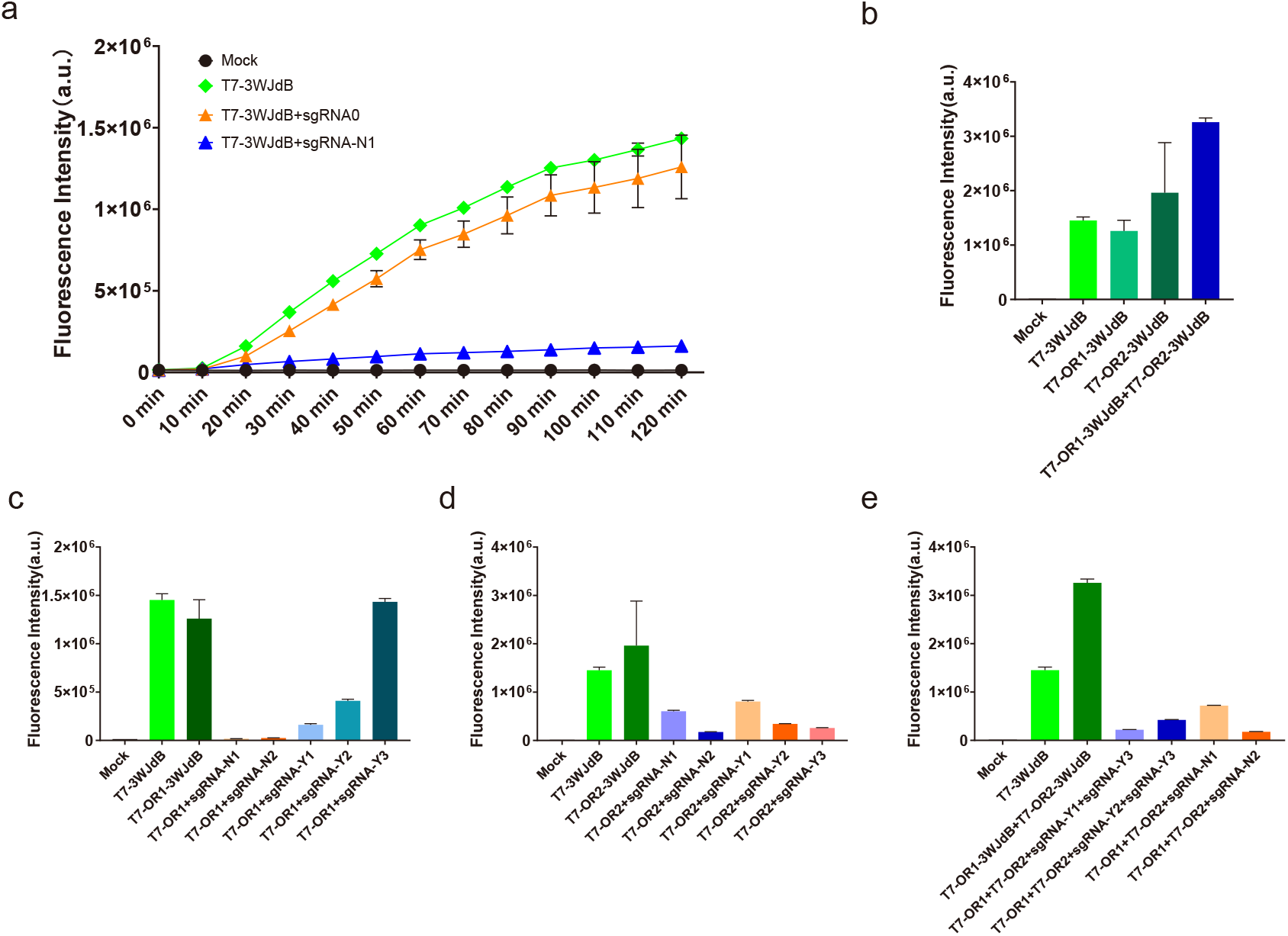
Real-time fluorescence data of CRISPRi inhibition reaction, where sgRNA-N1 targets a specific inhibition site and sgRNA0 is a nonspecific sgRNA. b: The effect of adding random orthogonal fragments on fluorescent aptamers. c and d: Inhibition of single-output reporter genes by different preset sgRNAs. e: Inhibition of dual-output reporter genes by different preset sgRNAs. T7-OR1-3WJdB is abbreviated as T7-OR1, and T7-OR2-3WJdB is abbreviated as T7-OR2.

### (3) Two-level dual-function Boolean logic gates

Studies have shown that adding sequences in the -35 promoter boxes does not interfere with the activity of an un-repressed promoter. Lei S.Qi et al. find targeting of the dCas9/sgRNA complex to the -35 box significantly knocked down gene expression (∼100 fold of repression)^43^. Research by Andriy Didovyk et al. demonstrated that by artificially designing and inserting random orthogonal fragments upstream of the -35 promoter boxes, a highly orthogonal set of dCas9/sgRNA complex/promoter pairs could be constructed. These dCas9/sgRNA complex/promoter pairs exhibit maximal orthogonality both among each other and with the host genome, addressing the issues of specificity and modularity in synthetic gene circuits.

Based on those, in the design of Boolean logic gates for CRISPRi construction, we propose a novel dual-function design focusing on two-level coupling. Specifically, we introduce “OR-NOT” (NOR) and “AND-NOT” (NAND) gates with a unified primary design strategy. Each gate consists of two distinct components, each comprising two different dsDNAs. Two different sgRNAs serve as inputs, and the outputs are a set of two different dsDNAs, both utilizing the 3WJdB fluorescent aptamer as the output signal (Fig. 3a). For NOR gate, either sgRNA-N1 or sgRNA-N2 can bind to the dCas9 protein within the system to inhibit fluorescence expression of the two output strands, thereby forming an NOR gate. Both output DNA strands are structurally identical, but each strand contains unique random orthogonal fragments (OR1 and OR2) that are completely different sequences, each with highly specific sgRNA sites (sgRNA-Y1 and sgRNA-Y3). Complete suppression of all fluorescent signals requires the simultaneous presence of both sgRNA-Y1 and sgRNA-Y3, constituting a logical NAND gate. All the related sequences can be found in Supplementary Materials. Before constructing NAND and NOR gates, we tested all the designed sgRNAs in the transcription system to choose the best sgRNAs to use (Fig. 2b-e).

**Figure 3:**
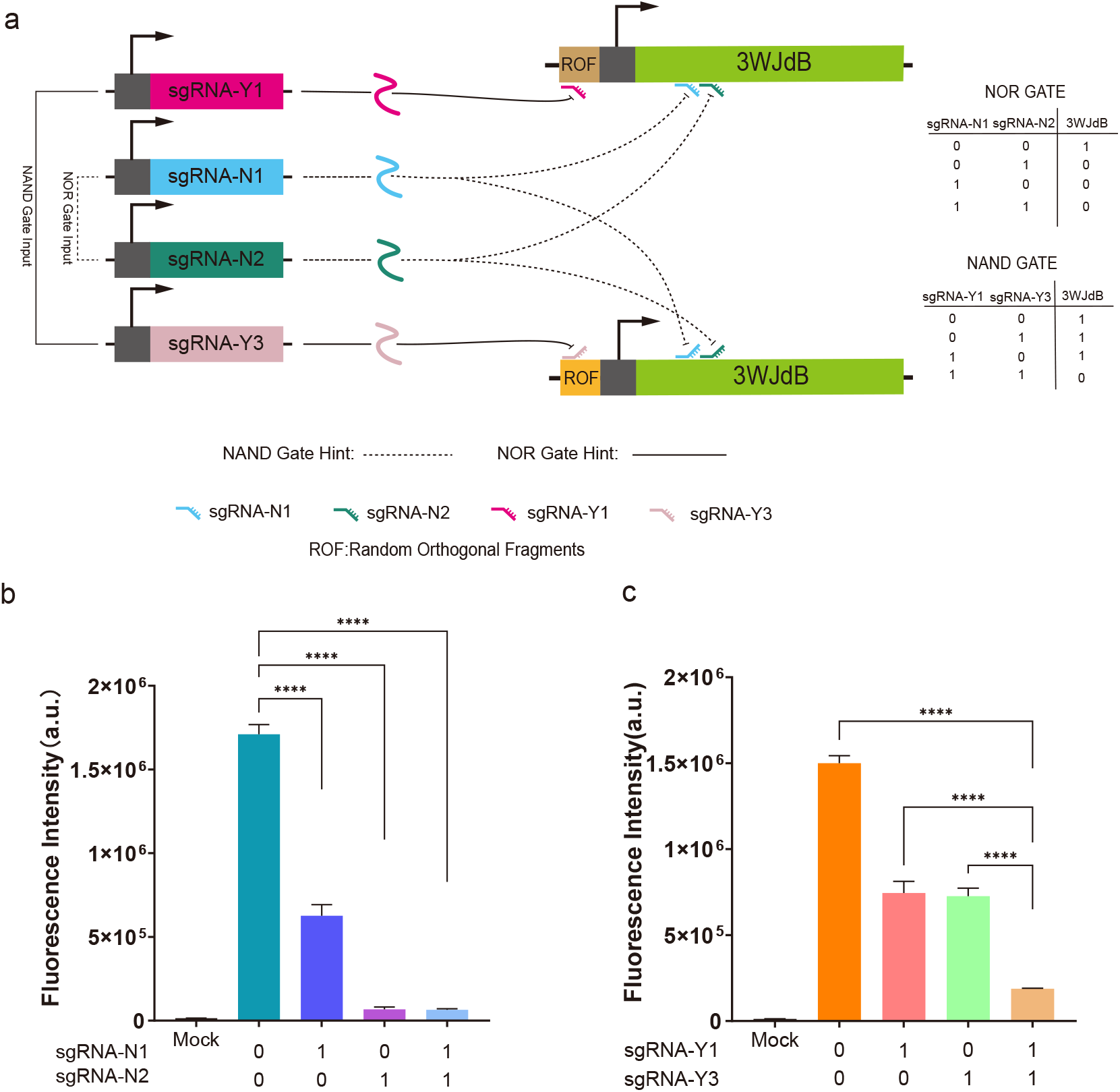
Schematic design of the NOR and NAND gates. The NOR gate is designed with inputs sgRNA-N1 and sgRNA-N2, where either input can inhibit the transcription of 3WJdB, thereby reducing the fluorescence signal. The NAND gate is designed with inputs sgRNA-Y1 and sgRNA-Y3, which target orthogonal random segments of the -35 region of the fluorescent aptamer promoter. Consequently, only the simultaneous presence of both sgRNAs can fully inhibit the fluorescence signal. b: Fluorescence output values of the NOR gate under different input conditions. c: Fluorescence output values of the NAND gate under different input conditions. Error bars represent the standard deviation (s.d.) of three biological replicates.

From the results of the NOR gate (Fig. 3b), we observed that expressing two CRISPRi nodes within the same T7 RNAP system did not present any abnormalities and maintained effective inhibition. When analyzing each CRISPRi node individually, it became evident that a single sgRNA targeting inhibition resulted in a weaker inhibitory effect compared to nodes with dual sgRNA targeting. This finding suggests that incorporating multiple sgRNA sites can enhance the inhibitory effect.

In the NAND gate design, each CRISPRi node functions independently. Consequently, the results (Fig. 3c) show that the fluorescence value, when only one node is inhibited, is nearly half of that observed when neither node is inhibited and is twice as high as when both nodes are inhibited. This indicates the independent and additive effects of the CRISPRi nodes on the output of the NAND gate.

### (4) Construction of Boolean Logic Gate by Three-Level Cascade

In Boolean algebra, a logical NOT operation can be seen as an “inverse,” as demonstrated in the NAND and NOR gate truth tables. NAND and NOR gates are essentially the inversions of AND and OR gates, respectively. Consequently, AND and OR gates can be derived by inverting the results of NAND and NOR gates. According to the law of double negation in Boolean algebra, any element remains unchanged after applying two NOT operations, expressed as (a’)’ = a. This means that inverting the results of NAND and NOR gates yields the AND and OR gates.

Building on this concept, we first tested whether CRISPRi nodes can be cascaded multiple times, and whether sgRNAs can be used as transmissible signals in biological circuits.

The logic of the three-level cascade can be summarized as follows: When the first-level sgRNA (sgRNA-T1-2 or sgRNA-T2-1) is introduced, it suppresses the output of the second-level sgRNA. As a result, the second-level sgRNA (sgRNA-T1 or sgRNA-T2) is inhibited, preventing the suppression of the final 3WJdB fluorescence signals, which leads to the generation of the overall fluorescence signal (Fig. 4a). Conversely, when the primary sgRNA is not introduced, the secondary sgRNA is produced normally. This secondary sgRNA then inhibits the final third-level containing 3WJdB gene, resulting in no overall fluorescence signal.

**Figure 4:**
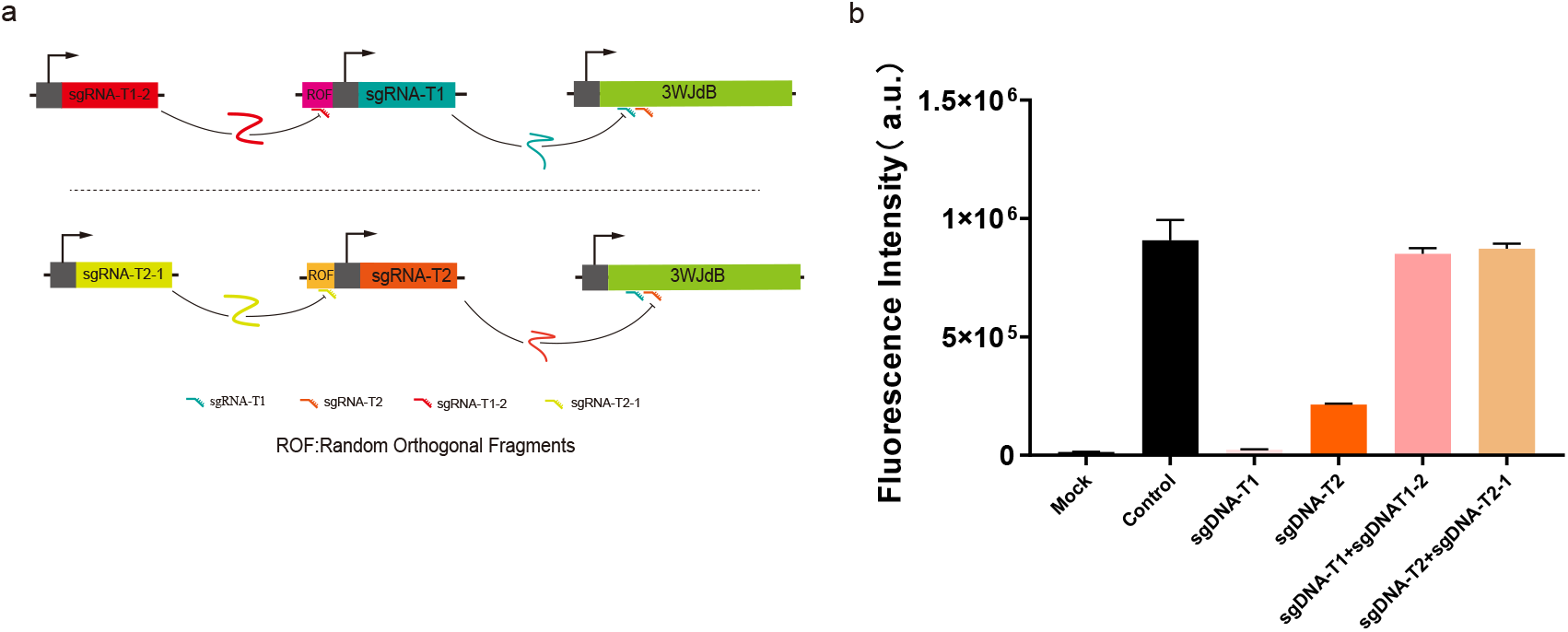
Schematic representation of a three-level cascade, where sgRNA functions as a signal that can be relayed multiple times. b: Fluorescence values of the three-level cascade. Secondary sgRNAs T1 and T2 can inhibit the transcription of the tertiary output fluorescent aptamer, while primary sgRNAs T1-2 and T2-1 can relieve the inhibition imposed by the secondary sgRNAs on the tertiary fluorescent aptamer. Error bars represent the standard deviation (s.d.) of three biological replicates.

For the third-level, we still used 3WJdB fluorescent aptamer as the output, and we designed the third-level sequence such that the third-level can be controlled by both the second-level sgRNA-T1 and sgRNAT-2, so that we can reuse the output module. As shown in Fig. 4b, sgRNA-T1 or sgRNA-T2 alone can suppress the transcription of the output module. And with the addition of the corresponding first-level sgRNAs, the suppression of the transcription can be reverted, indicating that sgRNA can be transmitted through multiple levels.

Then we designed a three-level AND and OR gates following the above-mentioned dual-function design. As depicted in the Fig. 5a, the designs of the AND and OR gates share most of the parts, differing only in their inputs. Both gates are structured into three levels. At the first level, the inputs are configured as: the AND gate uses the sgRNA-T1-2 and sgRNA-T2-1, which were selected for their optimal inhibitory efficiencies in a single three-level cascade validation experiment (Fig. 4). The OR gate uses two different inputs, sgRNA-S1 and sgRNA-S2, which are newly redesigned for the sgRNA handle sequence. Each sgRNA consists of a 20 nt specific binding sequence and a 60-80 nt hairpin structure that binds to the dCas9 protein, forming a complex.

**Figure 5:**
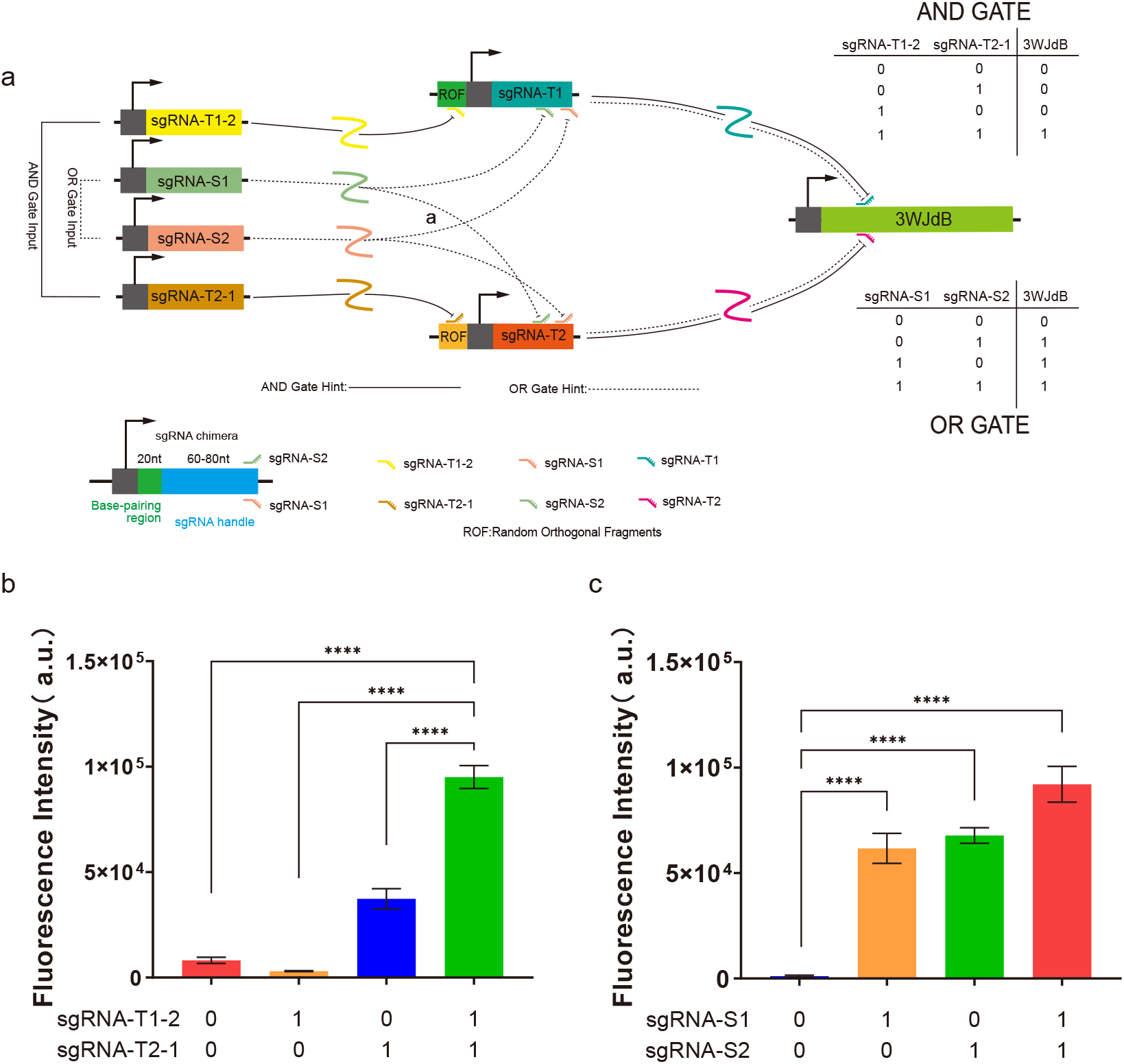
Schematic design of the AND and OR gates. The OR gate is formed with inputs sgRNA-S1 and sgRNA-S2, where either input can relieve the transcriptional inhibition of the secondary sgRNAs on the third-level output signal 3WJdB. The AND gate is formed with inputs sgRNA-T1-2 and sgRNA-T2-1, which target orthogonal random segments of the -35 region of the secondary sgRNA promoter. Consequently, only the simultaneous presence of both sgRNAs can fully relieve the inhibition on 3WJdB transcription. b: Fluorescence output values of the AND gate under different input conditions. c: Fluorescence output values of the OR gate under different input conditions. Error bars represent the standard deviation (s.d.) of three biological replicates.

As shown in Fig. 5a, both the AND and OR gates involve three CRISPRi nodes. For the AND gate, the two sgRNA-T nodes are designed to work in tandem to ensure that both must be inhibited for the final 3WJdB node to produce a signal. As we can see in Fig. 5b, the AND gate only showed a significant signal when both inputs were present, consistent with our expectation. Meanwhile, in the OR gate, the same two sgRNA-T nodes can be suppressed by either input. As shown in Fig. 5c, either one input or two inputs can turn the logic gate on to generate significant fluorescence signals, hence the OR gate. In summary, just like the NOR and NAND gates that we constructed previously, the three-level AND and OR gates utilize the same dual-function design, where by altering only the inputs, we can switch to different logic gates. These results reinforce the effectiveness of our modular approach in creating versatile and efficient synthetic gene circuits.

## Conclusions

Constructing synthetic gene circuits based on CRISPRi remains a promising alternative choice despite some challenges. Their versatility has been validated across various organisms, including bacteria^23,44,45^, mammalian cells^46–48^, yeast^11,49,50^, and even plant cells^18,51^. Complex gene circuits have been successfully constructed, demonstrating the wide applicability of CRISPRi in synthetic biology. Our work aims to enhance the integration and improve the coupling of synthetic gene circuits within host cells, streamlining the redundant segments of multiple gene circuits in the same environment.

In this study, we propose and validate a dual-function Boolean logic gate design approach that significantly reduces redundant design. Our design of NOR and NAND gates with identical information processing and output modules, only differing in their sgRNAs, demonstrates a significant reduction in workload and complexity. The verification of multi-level cascading of sgRNAs paves the way for the design of more sophisticated logic gates, such as multi-level AND and OR gates, which can be realized by inverting the results of the NAND and NOR gates.

Besides, when designing sgRNAs and ROFs, by performing bioinformatics analysis on the insertion fragments in the promoter -35 region, we ensure strict orthogonality among similar fragments, thereby reducing signal crosstalk^52^. And using fluorescent aptamers as reporter genes can further minimize the resource consumption by the entire synthetic gene circuit, reducing the potential impact on host cell growth.

Moving forward, we envision several next steps to further advance this field:

First, our main platform for this work is based on in vitro transcription system as it has been shown to be a valuable testbed for biological circuits. Next, we plan to test these logic gates in cells, further construct more complex gene circuits and eventually Biological Central Processing Unit (BCPU).

Second, in this work, we employed a dual-function design where logic gates share identical information processing and output modules, differing only in their sgRNA inputs. For instance, both the AND and OR gates utilize the same intermediate and output levels, but they have distinct inputs, each comprising two sgRNAs. By combining two specific sgRNAs, we can construct an AND gate, while another pair can generate an OR gate. Interestingly, with four input sgRNAs, there are six possible combinations, each potentially producing a unique output pattern—some of which may correspond to logic gates, while others may not. This diversity of potential outputs highlights the versatility and flexibility of our design approach, offering a broad spectrum of applications in synthetic biology and demonstrating the intricate possibilities that arise from such modular and combinatorial strategies.

## Acknowledgments

We would like to thank Dr. Xianai Shi, Dr. Jianmin Yang, Dr. Yunquan Zheng, Dr. Feng Li, Dr. Mingmao Chen, and Dr. Li Chen for helpful discussion and suggestions. This work was supported by the National Natural Science Foundation of China (32001037); Special project of Fujian Provincial Department of Finance (202309).

## Note

The authors declare no competing financial interest.

## Author contribution

Zhiwu Yao: conceptualization, data curation, formal analysis, investigation, methodology, visualization, original draft preparation. Shaobin Guo: conceptualization, project administration, resources, supervision, validation, original draft preparation and review and editing. All authors have read and agreed to the published version of the manuscript.

## Notes

### Competing Interest Statement

The authors have declared no competing interest.

